# Y chromosome damage underlies testicular abnormalities in ATR-X syndrome

**DOI:** 10.1101/2023.10.21.562414

**Authors:** Nayla Leon Carlos, Thanh Nha Uyen Le, Andrew Garvie, Lee H Wong, Stefan Bagheri-Fam, Vincent R Harley

**Affiliations:** Hudson Institute of Medical Research, Melbourne, Victoria 3168, Australia; Department of Molecular & Translational Science, Monash University, Melbourne, Victoria 3168, Australia; Monash University, Melbourne, Victoria 3800, Australia

**Keywords:** ATRX, ATR-X syndrome, DSD, testis, Sertoli cells, PML NB, GATA4 foci, Y-chromosome

## Abstract

ATR-X (<u>al</u>pha <u>t</u>halassemia, <u>m</u>ental <u>r</u>etardation, <u>X</u>-linked) syndrome is a severe developmental disorder affecting males caused by mutations in the chromatin remodelling gene *ATRX*. Genital abnormalities in affected boys include hypospadias and ambiguous genitalia, and patients show small poorly formed testes with only a few seminiferous tubules. Our mouse model recapitulated these testicular defects when *Atrx* was specifically deleted in Sertoli cells (Sc*Atrx*KO). Sc*Atrx*KO mice develop small testes with fewer and discontinuous tubules due to G2/M arrest and apoptosis of Sertoli cells.

Here, we investigated the mechanism underlying the Sertoli cell defects in ATR-X syndrome. In healthy male control mice, Sertoli cell nuclei contain a single novel “GATA4 PML nuclear body (NB)” that strongly expresses the transcription factor GATA4, as well as ATRX and its binding partner DAXX. The GATA4 PML NB co-localizes with heterochromatin protein HP1α and PH3 (a marker of chromosome condensation), and with the short arm of the Y chromosome (Yp). In contrast, Sc*Atrx*KO Sertoli cells contain a single giant GATA4 PML NB, frequently associated with DNA double-strand breaks in G2/M-arrested Sertoli cells that underwent apoptosis. HP1α and PH3 were absent from the giant GATA4 foci suggesting a local failure in heterochromatin formation and chromosome condensation. Our data indicate that in Sertoli cells, ATRX protects a chromosomal region of Yp from DNA damage, probably during replication stress, and thus protects Sertoli cells from cell death. We discuss Y chromosome damage as a novel mechanism for testicular failure and the potential role of GATA4 during this process.

**Disclosure Summary:** The authors have nothing to disclose.

## Introduction

ATRX (alpha thalassemia, mental retardation, X-linked) is a chromatin remodelling protein which belongs to the switch/sucrose non-fermentable (SWI-SNF) family. Mutations in *ATRX* cause the ATRX-syndrome, which is characterized by distinct craniofacial features, severe intellectual disability, alpha thalassemia, and urogenital abnormalities (Gibbons & Higgs, 2000; Gibbons et al., 1991; Wilkie et al., 1991; Wilkie et al., 1990). ATRX has multiple important biological functions such as in the formation and maintenance of pericentromeric and telomeric heterochromatin, chromosome condensation, and in the protection of common fragile sites, pericentromeric heterochromatin, and telomeres during replication (Pladevall-Morera et al., 2019). ATRX can recruit Heterochromatin Protein 1 alpha (HP1α) at pericentromeric heterochromatin (Marano et al., 2019). Within PML nuclear bodies (PML NBs) ATRX and its interacting partner Death Domain Associated Protein (DAXX) deposit the histone variant H3.3 at pericentromeric heterochromatin and telomeres (Drané et al., 2010; Goldberg et al., 2010; Tang et al., 2004; Wong et al., 2010; Xue et al., 2003). During replication, ATRX and DAXX facilitate replication by preventing G4 quarter DNA structure formation and promote replication fork recovery and homologous recombination-dependent repair of DNA double strand breaks (Clynes et al., 2014; Juhász et al., 2018; Leung et al., 2013; Pladevall-Morera et al., 2019; Raghunandan et al., 2020; Watson et al., 2013; Wong et al., 2010).

PML NBs are sub-nuclear structures involved in various cellular functions such as replication, gene regulation, apoptosis, heterochromatin condensation, telomere integrity and DNA damage response (Bernardi & Pandolfi, 2007; Bernardi et al., 2008; Chang et al., 2018; Dellaire et al., 2006; Lallemand-Breitenbach & de Thé, 2010; Salomoni & Pandolfi, 2002; Takahashi et al., 2004). They are extensively distributed, being found in most cell-lines and many tissues. The number of PML NBs varies from 5 to 30 per nucleus. Under normal conditions, PML NBs size range from 0.1 to 1.0 µm in diameter (Melnick et al., 1999; Weis et al., 1994). The protein components of the inner core are highly variable. To date, over 250 different proteins have been described to be recruited by PML NBs including ATRX and DAXX (Barroso-Gomila et al., 2021; Dellaire et al., 2003; Van Damme et al., 2010). Recently, the short arm of the Y chromosome has been identified as a region to which PML NBs frequently bind (Kurihara et al., 2020). No function of PML NBs during sex development or in differences/disorders of sex development (DSD) has been demonstrated to date.

A common co-morbidity in ATR-X syndrome is a difference of sex development (DSD) affecting XY individuals. Patients display genital abnormalities varying in severity from cryptorchidism to complete female external genitalia. Gonadal histology reveals small testes containing only a few seminiferous tubules (Ion et al., 1996; Wilkie et al., 1990). Previously we generated an *Atrx* knockout mouse model with *Atrx* specifically inactivated in Sertoli cells (Sc*Atrx*KO), a cell lineage crucial for formation of testis cords, the presumptive seminiferous tubules. Like boys with ATR-X syndrome, Sc*Atrx*KO mice showed small testes with fewer tubules. The lack of tubules was due to G2/M cell cycle arrest and apoptosis of Sertoli cells during foetal life (Bagheri-Fam et al., 2011).

In this study, we describe a single novel PML nuclear body we call “GATA4 PML NB” in testicular Sertoli cells that encapsulates the short arm of the Y chromosome and expresses the transcription factor GATA4. The GATA4 PML NB contains ATRX, DAXX, HP1α and shows early enrichment of PH3, a marker of chromosome condensation at G2. In Sc*Atrx*KO mice, these GATA4 PML NBs are enlarged and are associated with DNA double stranded breaks in G2/M-arrested Sertoli cells that undergo apoptosis. The absence of HP1α and PH3 at the GATA4 foci in giant PML NBs indicate a defect in heterochromatin formation and chromosome condensation, respectively. Our results suggest Y chromosome damage as a novel mechanism in the aetiology of ATRX-syndrome.

## RESULTS

### A single novel PML NB is enlarged in *Atrx* knockout Sertoli cells

Conditional knockout of *Atrx* in the Sertoli cells of the fetal mouse testis (Sc*Atrx*KO) leads to cell cycle arrest at G2/M and associated apoptosis (Bagheri-Fam et al., 2011). With further investigation, we observed an unusual staining pattern of GATA4 in Sertoli cells of both control and Sc*Atrx*KO gonads (**Fig.1A**). GATA4 is important for testis development and reportedly shows diffuse expression throughout the nucleus (Manuylov et al., 2011; Tevosian et al., 2002). In addition to this diffuse expression, we found that around 10% of control and 26% of Sc*Atrx*KO Sertoli cells contain a single GATA4 immunoreactive nuclear speckle (GATA4 foci), frequently located near the periphery of the nucleus (**Fig. 1A and Table 1**). Some GATA4 foci were significantly larger in Sc*Atrx*KO when compared to control testes; the size range in the control testes was 0.1-1µm, whereas in the Sc*Atrx*KO it was 0.2-2µm, with around 29% of Sertoli cells containing enlarged (giant) GATA4 foci outside of the control range (>1.0µm; **Fig. 1E**).

**Figure 1.**
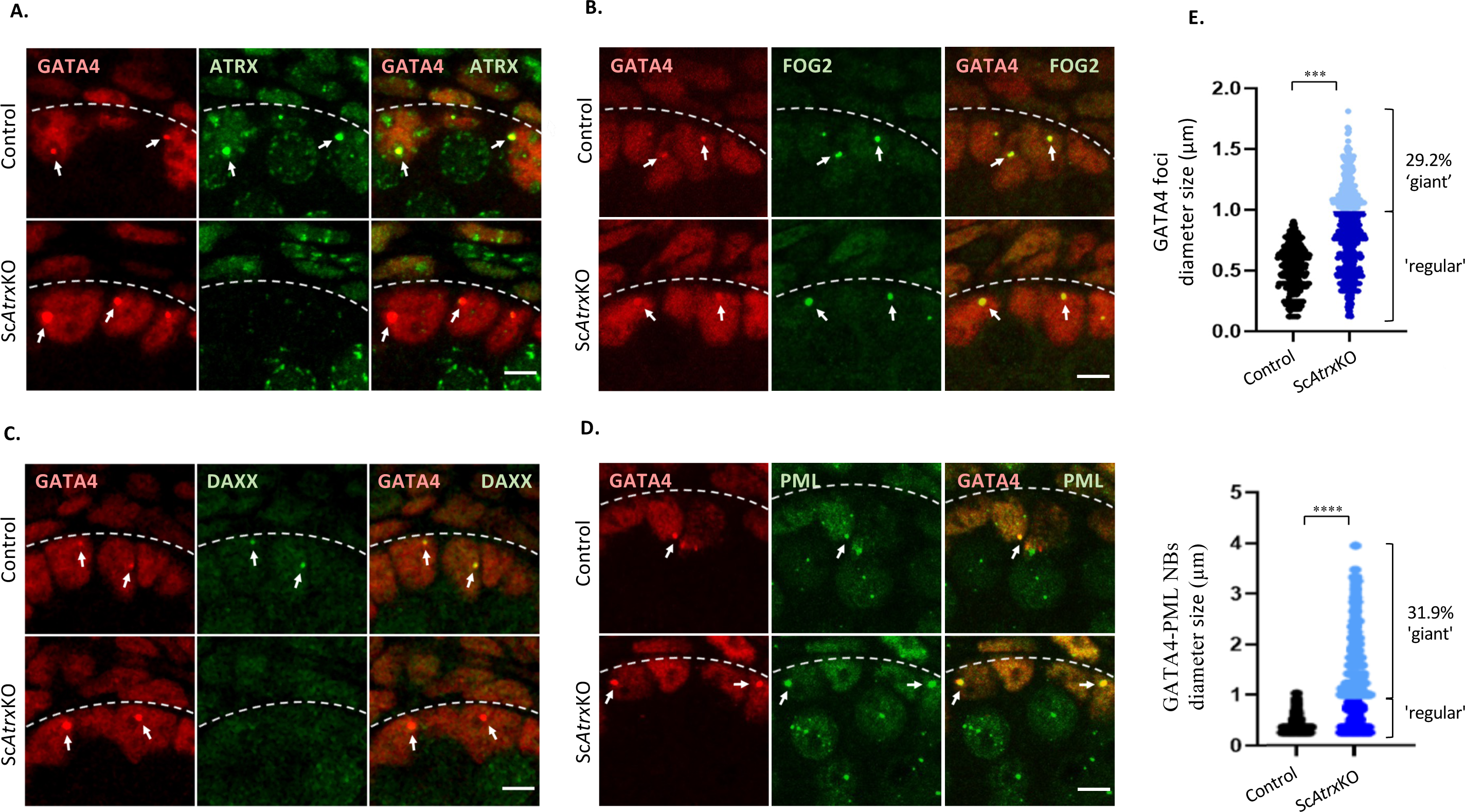
Sertoli cells contain a novel GATA4 PML NB which is enlarged in the absence of ATRX. (**A-D**) Double immunofluorescence (IF) analyses in E16.5 XY control and Sc*Atrx*KO testes. Dashed lines mark the basal lamina of the testis cords. White arrows denote GATA4 foci. At least 100 GATA4 foci were analysed per testis (n=3). Scale bars are 5μm. (**A**) Double IF for GATA4 (red, nuclear) and ATRX (green, nuclear). (**B**) Double IF for GATA4 (red, nuclear) and FOG2 (green, nuclear). (**C**) Double IF for GATA4 (red, nuclear) and DAXX (green, nuclear). (**D**) Double IF for GATA4 (red, nuclear) and PML (green, nuclear). (**E**) Measurements of GATA4 foci and GATA4-PML NB diameters. More than 100 GATA4 foci and GATA4-PML NBs were measured in each testis (control, n=3; Sc*Atrx*KO, n=4), using FUJI software. The control is represented in black and the Sc*Atrx*KO in blue. Dark blue shows GATA4 foci and GATA4-PML NBs that have a regular size in Sc*Atrx*KO Sertoli cells, while light blue shows GATA4 foci and GATA4-PML NBs outside the control range. ***p< 0.001, ****p< 0.0001; unpaired two-tailed t-test compared to control.

**Table 1.** Percentage of Sertoli cells with single GATA4 foci in control and Sc*Atrx*KO testes.

|  | **Control** | **KO** |
| --- | --- | --- |
| Animals | n=3 | n=3 |
| Number of Sertoli cells | 1145 | 962 |
| Sertoli cells with single GATA4 foci | 119 | 251 |
| **Percentage of Sertoli cells with single GATA4 foci** | **10.4%** | **26.1%** |

To investigate if these giant GATA4 foci could be a direct consequence of ATRX loss at these sites, we performed co-immunofluorescence (IF) for GATA4 and ATRX (**Fig. 1A and Fig. S1**). As expected, ATRX showed diffuse nuclear expression, and moderate levels at the bright DAPI regions (chromocenters) that contain the highly compacted pericentromeric heterochromatin (major satellites) of the chromosomes (Guenatri et al., 2004). Sertoli cells also showed a single nuclear speckle with high ATRX IF staining intensity that co-localized with the GATA4 foci, located near one of the bright DAPI regions (**Fig. 1A and Fig. S1**). This suggests that ATRX has an important function at these foci. We next looked for the presence of two well-studied interacting partners of GATA4 and of ATRX at the GATA4 foci, FOG2 and DAXX, respectively. Co-IF for GATA4 and FOG2 revealed co-localization in both control and Sc*Atrx*KO Sertoli cells (**Fig. 1B**). While DAXX co-localized with the GATA4 foci in control gonads, it was absent in Sc*Atrx*KO Sertoli cells (**Fig. 1C**), indicating that presence of DAXX depends on ATRX.

ATRX and DAXX are well-known components of PML nuclear bodies (PML NB). We investigated whether GATA4 foci might associate with PML NBs. Co-IF for GATA4 and PML in control Sertoli cells revealed that 17% of GATA4 foci co-localized with one specific PML NB, whereas the other 83% were PML-negative (**Fig. 1D and Table 2**). In contrast, in Sc*Atrx*KO testes, 77% of the regular sized GATA4 foci (up to 1.0µm) co-localized with the PML NB and almost all (94%) of the giant GATA4 foci (>1.0µm) were positive for PML. Around 32% of GATA4 PML NBs in Sc*Atrx*KO Sertoli cells were larger than the control range (**Fig.1E**). Therefore, the giant GATA4 foci are the giant PML NBs in the Sc*Atrx*KO Sertoli cells. Taken together, these data suggest that PML is recruited to the ATRX-GATA4 foci in both control and Sc*Atrx*KO Sertoli cells, but to far a greater extent in Sc*Atrx*KO Sertoli cells.

**Table 2.** Percentage of PML-positive GATA4 foci in control and Sc*Atrx*KO testes.

|  | **Control** | **KO** |  |
| --- | --- | --- | --- |
| Animals | n=3 | n=3 |  |
| Number of Sertoli cells with single GATA4 foci | 119 | 216 (regular sized) | 35 (giant) |
| PML-positive GATA4 foci | 20 | 166 (regular sized) | 33 (giant) |
| **Percentage of PML-positive GATA4 foci** | **16.8%** | **76.9% (regular sized)** | **94.3% (giant)** |

### Those Sc*Atrx*KO Sertoli cells with giant GATA4 PML NBs are arrested at G2/M

The G2/M phase of the cell cycle is prolonged in Sertoli cells of Sc*Atrx*KO mice when compared to control mice (Bagheri-Fam et al., 2011). However, the underlying mechanism remained elusive. We examined if the giant GATA4 foci in Sc*Atrx*KO Sertoli cells are related to the cell cycle defect, by co-IF for GATA4 and for phospho-histone H3 (PH3), a cell cycle marker of late G2 phase and mitosis. (Hendzel et al., 1997). PH3 IF staining confirmed that Sc*Atrx*KO testes have more Sertoli cells at G2/M than do control testes (23.5% versus 14.6%) (**Fig. 2B**). When considering only those Sertoli cells with GATA4 foci, control testes have more GATA4 foci positive Sertoli cells at G2/M (49.6%) when compared to the entire Sertoli cell population (14.6%) (**Fig. 2B**). Similarly, the percentage of Sc*Atrx*KO Sertoli cells with regular sized GATA4 foci at G2/M (44.6%) (**Fig. 2C**) was higher than in the entire Sertoli cell population (23.5%) (**Fig. 2B**). Strikingly, Sc*Atrx*KO Sertoli cells with giant GATA4 foci were all at G2/M, showing strong speckled PH3 staining typical of cells at late G2 phase (**Fig. 2A and C**) (Hendzel et al., 1997).

**Figure 2.**
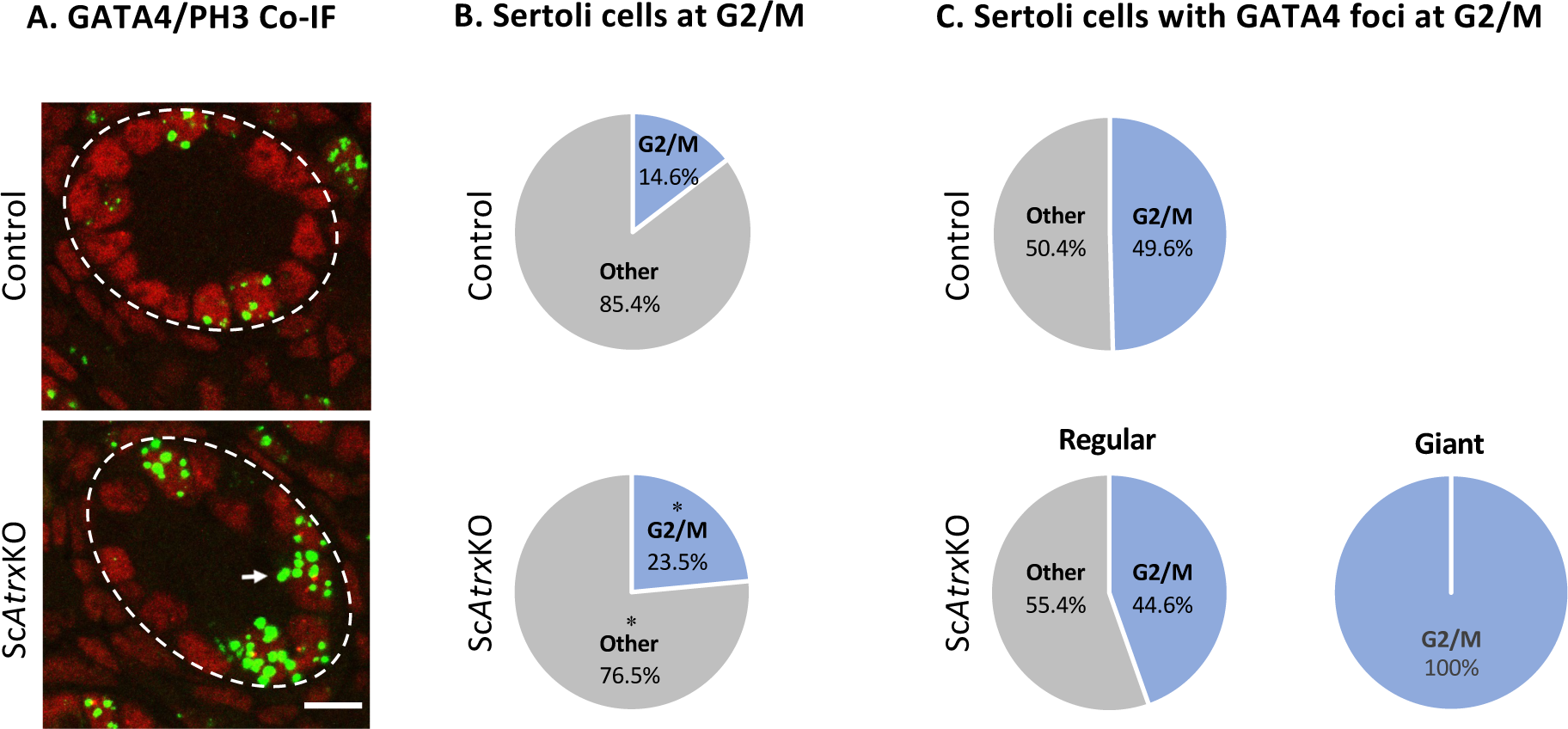
Sc*Atrx*KO Sertoli cells with giant GATA4 foci are arrested at G2/M. (**A**) Double immunofluorescence (IF) analyses in E16.5 XY control and Sc*Atrx*KO testes for GATA4 (red, nuclear) and PH3 (green, nuclear), a marker of late G2 phase and mitosis. Dashed lines mark the basal lamina of the testis cords. The white arrow denotes a Sertoli cell with a giant GATA4 speckle. Scale bar is 10μm. (**B**) Pie charts showing the percentages of the cell cycle stages in Sertoli cells of E16.5 XY control and Sc*Atrx*KO testes. More than 2000 and 1000 Sertoli cells were counted in n=2 control and n=3 Sc*Atrx*KO testes, respectively. *p< 0.05, one-way ANOVA. (**C**) Pie charts showing the percentages of the cell cycle stages in Sertoli cells with GATA4 foci of E16.5 XY control and Sc*Atrx*KO testes. More than 200 Sertoli cells with GATA4 foci were analysed in n=2 control and n=3 Sc*Atrx*KO testes.

These data show that Sertoli cells with GATA4 foci in both control and Sc*Atrx*KO testes are enriched at late G2/M phase. Furthermore, all Sc*Atrx*KO Sertoli cells with giant GATA4 foci are arrested at G2/M. Therefore, it is likely that a defect at the enlarged GATA4 foci underlies the G2/M cell cycle arrest in Sc*Atrx*KO Sertoli cells.

### Loss of ATRX at the GATA4 foci leads to DNA double strand breaks and Sertoli cell apoptosis

ATRX reportedly protects common fragile sites, pericentromeric heterochromatin and telomeres prone to DNA double-strand breaks (DSBs) under conditions of replication stress. DSBs trigger the cell to activate the DNA damage response to repair DNA (Plesca et al., 2008). One of the earliest and important chromatin changes during this process is the phosphorylation of the Histone variant H2AX (γ-H2AX) around the sites of the DNA double-strand breaks (Rogakou et al., 2000). γ-H2AX recruits DNA repair proteins and induces G2/M cell cycle arrest which will result in either DNA repair or apoptosis. γ-H2AX enrichment also occurs as a response to the DNA fragmentation during apoptosis (Rogakou et al., 2000).

To investigate whether DSBs underlie the G2/M arrest and apoptosis of Sc*Atrx*KO Sertoli cells, we performed co-IF for GATA4 and γ-H2AX (**Fig. 3A)**. In both control and Sc*Atrx*KO Sertoli cells, we observed two different types of γ-H2AX staining patterns in Sertoli cells with GATA4 foci. γ-H2AX showed either localized staining (L) at the GATA4 foci only (**Fig. 3A and B**, blue slices in pie charts), or both localized and widespread (LW) throughout the nucleus, indicative of the cell undergoing apoptosis (**Fig. 3A and B**, orange slices in pie charts). In control testes, around 10% of the Sertoli cells with GATA4 foci showed prominent γ-H2AX staining (4.1% localized and 6.2% widespread) (**Fig. 3B)**. Sc*Atrx*KO Sertoli cells with regular sized GATA4 foci showed similar widespread γ-H2AX staining (5.8%), but localized γ-H2AX staining was increased by almost 11-fold (43.7%) (**Fig. 3B)**. Remarkably, all G2/M-arrested Sertoli cells with the giant GATA4 foci were γ-H2AX-positive with 51.7% and 48.3% of them showing localized and widespread γ-H2AX-staining, respectively (**Fig. 3B)**. To confirm the presence of DSB at the GATA4 foci we also performed co-IF for the chromatin-binding p53-binding protein 1 (53BP1) which is an important regulator of DSB signalling by recruiting DSB signalling and repair proteins to the damaged site and promoting end-joining of distal DNA ends (Panier & Boulton, 2014). Indeed, while 53BP1 staining was present at the GATA4 foci in Sertoli cells of Sc*Atrx*KO testes, it was not detectable in control testes (**Fig. S2**).

**Figure 3.**
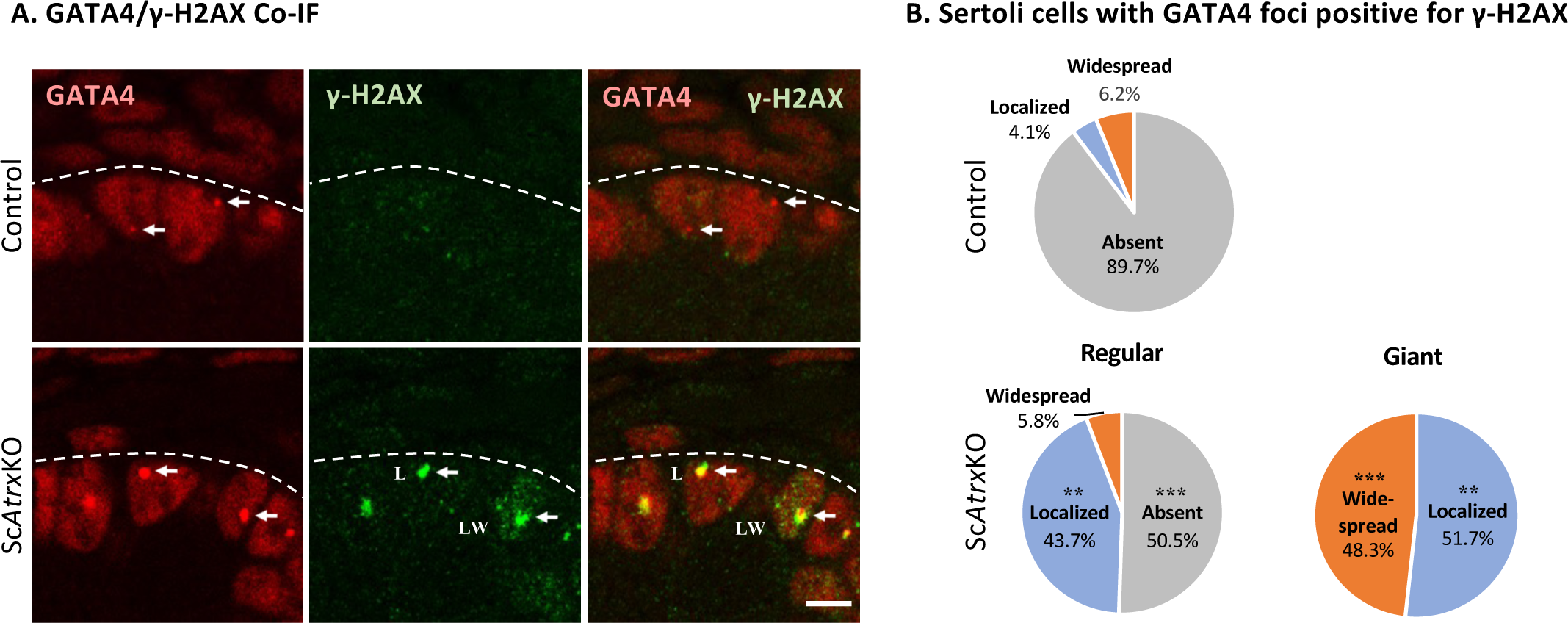
Loss of ATRX at the GATA4 foci leads to DNA double strand breaks and apoptosis. (**A**) Double immunofluorescence (IF) analyses in E16.5 XY control and Sc*Atrx*KO testes for GATA4 (red, nuclear) and γ-H2AX (green, nuclear), a marker of DNA double strand breaks and apoptosis. White arrows denote GATA4 foci. L, localized γ-H2AX staining specifically at the GATA4 foci; LW, localized and widespread γ-H2AX staining throughout the nucleus, indicative of the Sertoli cell undergoing apoptosis. Dashed lines mark the basal lamina of the testis cords. Scale bar is 5μm. (**B**) Pie charts showing the percentages of γ-H2AX-positive GATA4 foci in Sertoli cells of E16.5 XY control and Sc*Atrx*KO testes. More than 200 Sertoli cells with GATA4 foci were analysed in n=2 control and n=3 Sc*Atrx*KO testes. **p<0.01, ***p<0.000; one-way ANOVA.

Taken together, these data indicate that loss of ATRX at the GATA4 foci leads to local DNA damage followed by apoptosis of the G2/M-arrested Sertoli cells with giant GATA4 foci.

### GATA4 foci represent sites of early chromosome condensation that require ATRX

Phosphorylation of histone 3 is required for the initiation of chromosome condensation (Van Hooser et al., 1998). H3 phosphorylation initiates at the pericentromeric heterochromatin of the chromosomes (chromocenters, bright DAPI regions) during G2 phase, then spreads towards the chromosome arms and is complete by mitotic prophase (Hendzel et al., 1997). Surprisingly, our co-IF analyses for GATA4 and PH3 revealed that PH3 staining was not only found at the bright DAPI regions, but also at the GATA4 foci (**Fig. 4A**). In PH3-positive control Sertoli cells, 86% of GATA4 foci were PH3-positive (**Fig. 4B**). However, only 52% of the regular sized GATA4 foci in PH3-positive Sc*Atrx*KO Sertoli cells were also positive for PH3, whereas no PH3 staining was found at the giant GATA4 foci in G2/M-arrested Sertoli cells (**Fig. 4B**). In contrast, H3 phosphorylation at the bright DAPI regions in Sc*Atrx*KO Sertoli cells was unaffected (**Fig. 4A**, yellow arrowheads). These data suggest that like the chromocenters, the GATA4 foci might represent sites of early chromosome condensation. They also suggest that loss of ATRX in Sertoli cells results in a failure of chromosome condensation at the GATA4 foci, but not at the chromocenters.

**Figure 4.**
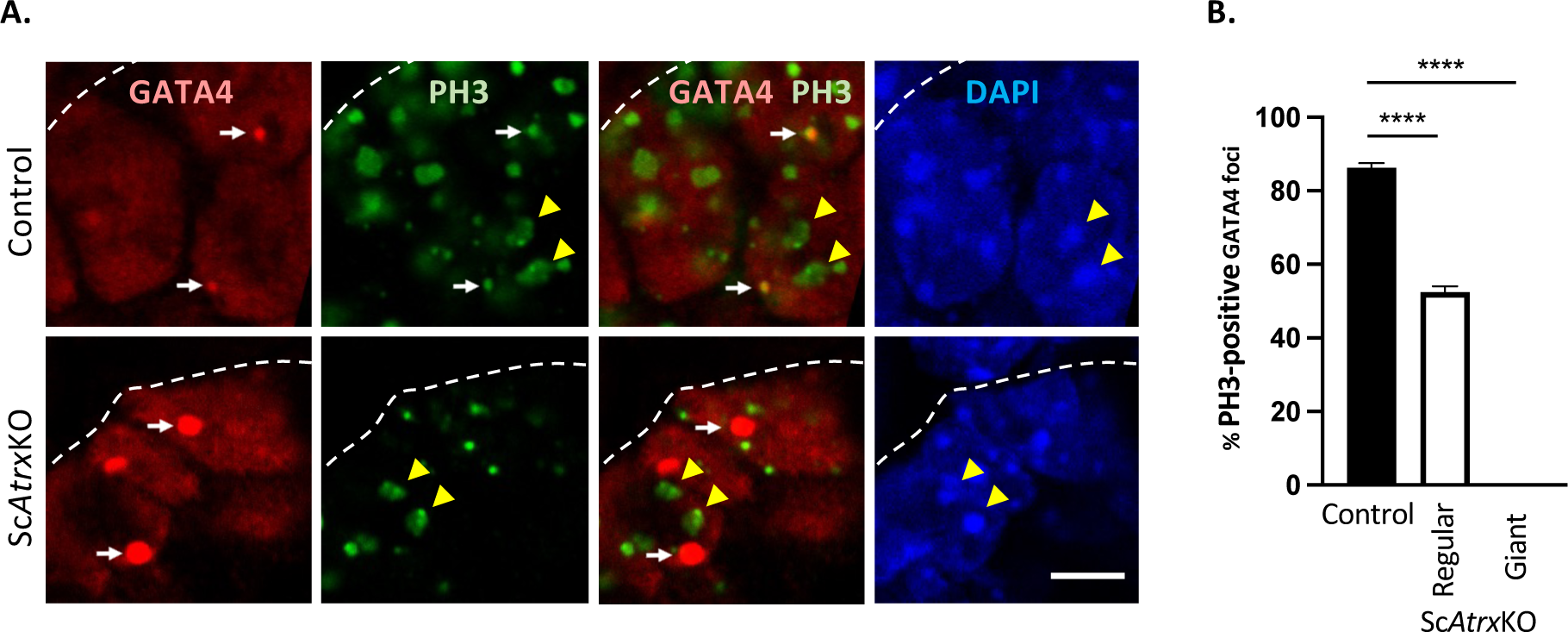
GATA4 foci are a site of early chromosome condensation that requir*es ATRX.* (**A**) Double immunofluorescence (IF) analyses in E16.5 XY control and Sc*Atrx*KO testes for GATA4 (red, nuclear) and PH3 (green, nuclear), a marker of late G2 phase and mitosis. DAPI (blue) was used as a nuclear stain. White arrows show GATA4 foci; yellow arrowheads denote PH3-positive pericentromeric heterochromatin (bright DAPI regions); Dashed lines mark the basal lamina of the testis cords; Scale bar is 5μm. (**B**) Percentage of PH3-positive GATA4 foci in Sertoli cells of E16.5 XY control and Sc*Atrx*KO testes. More than 200 Sertoli cells with GATA4 foci were analysed in n=2 control and n=3 Sc*Atrx*KO testes. Values are the mean ±SEM. ****p<0.0001; One-way ANOVA.

We next looked for potential structural similarities between the GATA4 foci and chromocenters. A characteristic feature of each chromocenter is the presence of highly compacted HP1⍺-positive pericentromeric heterochromatin (major satellites) of multiple chromosomes, while the corresponding centromeres (minor satellites) closely surround the chromocenters as several separate structures (Guenatri et al., 2004). Indeed, co-IF analyses in control Sertoli cells for GATA4 and HP1⍺, a hallmark of pericentromeric heterochromatin, revealed that 79% of GATA4 foci are HP1⍺-positive (**Fig. 5**). This suggests that HP1⍺ is recruited to the GATA4 foci and that the GATA4 foci, like the chromocenters, contain highly compacted heterochromatin. However, in Sc*Atrx*KO testes, HP1⍺ staining was either severely reduced or absent at the GATA4 foci. (**Fig. 5**), suggesting that heterochromatin formation at these foci is perturbed in the absence of ATRX. In contrast, HP1⍺ staining at pericentromeric heterochromatin (bright DAPI regions) in Sc*Atrx*KO testes was unaffected (**Fig. 5**, yellow arrowheads). Co-IF for GATA4 and the centromeric protein CENPA, which is found at all centromeres, revealed a single signal in close association with the GATA4 foci in both control and Sc*Atrx*KO Sertoli cells (**Fig. 6A**, left images). Furthermore, Co-IF for GATA4 and TERF1, a telomere marker, revealed two telomere signals closely associated with the GATA4 foci of control and Sc*Atrx*KO Sertoli cells (**Fig. 6A**, right images).

**Figure 5.**
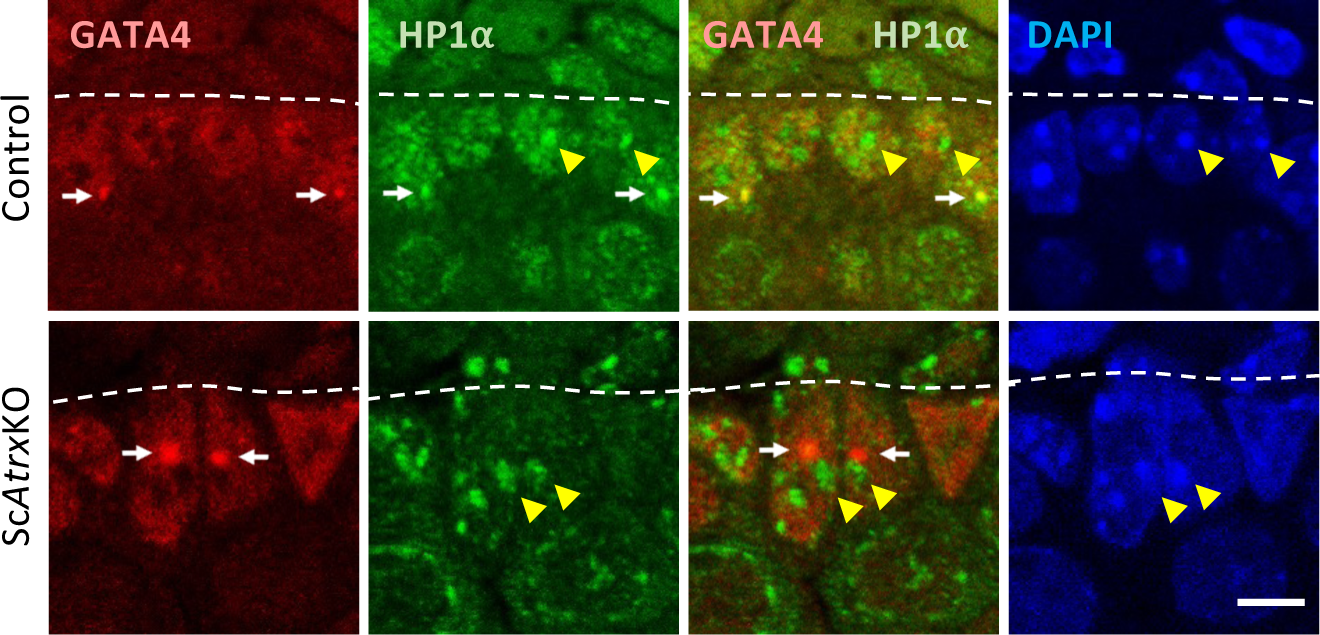
Sc*Atrx*KO Sertoli cells show specific loss of HP1⍺ at the GATA4 foci. Double immunofluorescence (IF) analyses in E16.5 XY control and Sc*Atrx*KO testes for GATA4 (red, nuclear) and HP1⍺ (green, nuclear), a marker of highly compacted heterochromatin. DAPI (blue) was used as a nuclear stain. White arrows show GATA4 foci; yellow arrowheads denote HP1⍺-positive pericentromeric heterochromatin (bright DAPI regions); Dashed lines mark the basal lamina of the testis cords. Scale bar is 5μm.

**Figure 6.**
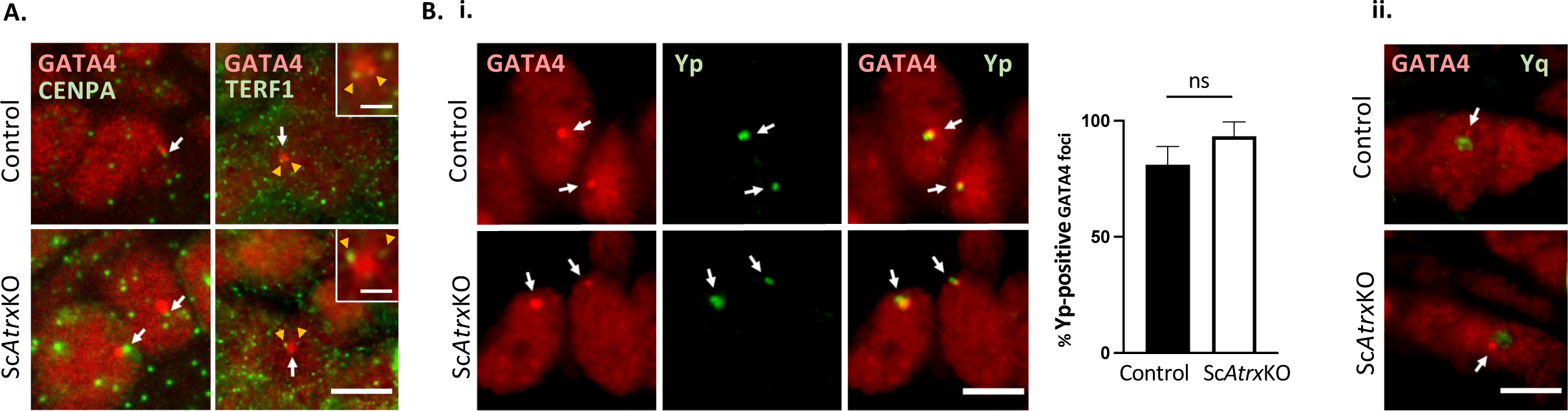
GATA4 foci contain the Y chromosome short arm (Yp). (**A**) *Left images:* Double immunofluorescence (IF) analyses in E16.5 XY control and Sc*Atrx*KO testes for GATA4 (red, nuclear) and CENPA (green, nuclear), a centromere marker. *Right images*: Double IF analyses in E16.5 XY control and Sc*Atrx*KO testes for GATA4 (red, nuclear) and TERF1 (green, nuclear), a telomere marker. The insets show magnifications of the GATA4 foci (**B**) **i**) Immuno-FISH for GATA4 and a Yp probe. Yp co-localized completely or partially with the GATA4 signal in 81% of control and 93% of Sc*Atrx*KO Sertoli cells. More than 150 Sertoli cells with GATA4 foci were analysed in control and Sc*Atrx*KO testes Values are the mean ±SEM (n=3 gonads). Ns; not significant; two tailed t-test compared to control. **ii**) Immuno-FISH for GATA4 and a Yq probe. Scale bars are 5μm; Scale bars in insets are 1μm.

Taken together, the GATA4 foci show structural and functional similarities to chromocenters (Presence of HP1⍺, PH3 and centromere in close association), but unlike chromocenters they appear to contain only chromatin from a single chromosome.

### GATA4 foci contain the Y chromosome short arm (Yp)

Recently, it has been demonstrated that a 300 kb region (YS300) on the short arm of the Y chromosome (Yp) is frequently associated with a large PML NB in mouse embryonic stem cells (Kurihara et al., 2020).Therefore, we investigated whether the single chromosome at the GATA4 foci may be the Y chromosome. We performed Immuno-FISH for GATA4 and two different Y probes, a 73.6 kb Yp probe (Chr Y: 112,036-184,662) that detects the 300 kb region associated with the reported PML NB (Kurihara et al., 2020) and a Yq probe that detects a 1.8 kb repeat dispersed over a 48 Mb region across the long arm of the Y chromosome (Chr Y: 4,173,195-52,194,425). Immuno-FISH for GATA4 and the Yp probe revealed that Yp co-localized completely or partially with the GATA4 signal in 81% of control and 93% of Sc*Atrx*KO Sertoli cells (**Fig. 6B**). In contrast, the signal of the Yq probe was often close to but did not-co-localize with the GATA4 signal (**Fig. 6C**). Taken together, these data indicate that the GATA4 foci locate to the short arm of the Y chromosome.

### Yp is enriched for GATA repeats and contains potential G4 DNA structures

Given the colocalization of the GATA4 foci with Yp, we wondered whether the Yp probe sequence contains potential ATRX and GATA4 binding sites. The GATA4 foci show intense staining, suggesting that large numbers of GATA4 proteins are present at these sites. It is therefore possible that GATA4 is binding repetitively to chromatin, unlike in its role as a transcription factor where it binds only once (Bouchard et al., 2019). The GATA family binds to the sequence motif ‘T/A(GATA)A/G’ with AGATAG being the most preferred (Merika & Orkin, 1993). Sex chromosomes contain highly conserved banded krait minor (Bkm) satellite DNA sequences in which GATA repeats are the major component. Intriguingly, in the mouse, these GATA repeats are predominantly confined to the short arm of the Y chromosome (Singh & Jones, 1982; Singh et al., 1994). Analysis of the entire 73.6 kb Yp probe sequence revealed a ∼4.6 kb region with a high accumulation of GATA repeats of which the most abundant is the preferred GATA4 binding site ‘AGATAG’ (**Fig. 7**). In contrast, the weaker GATA family binding sites ‘C/G(GATA)T/C’ are present in low numbers and are evenly distributed throughout the Yp sequence (**Fig. 7**). ATRX binds to G-quadruplex DNA secondary structures (G4) that are formed by G-rich tandem repeats such as the telomeric repeat TTAGGG (Wang & Patel, 1992). The G4 DNA sequence consist of four interrupted stretches of at least three guanines which form a four-stranded DNA secondary structure that is stabilized by G-quartets. Using QGRS Mapper (Kikin et al., 2006), we identified 8 putative G4□sequences with a G-score threshold ≥ 22 (**Table 3**). These data show that the 73.6 kb Yp probe sequence contains potential binding sites for both GATA4 and ATRX.

**Figure 7.**
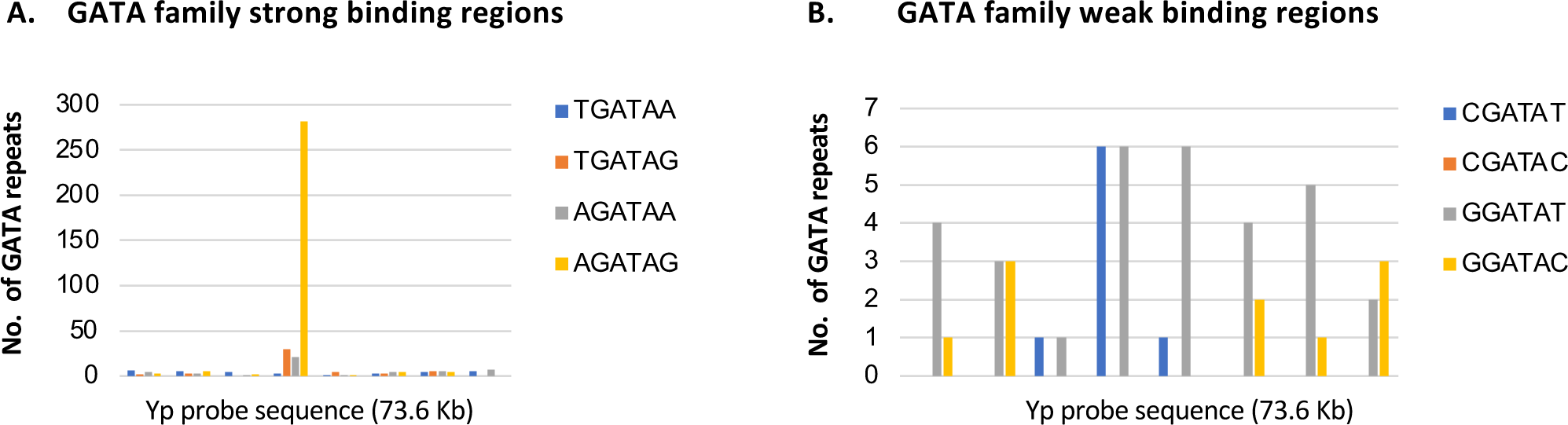
Yp is enriched for GATA repeats. (**A**) The 73.6 kb Yp probe contains a 4.6. kb region that is enriched for the strong GATA family binding site ‘AGATAG’. (**B**) The GATA family poor binding sites, ‘C/G(GATA)T/C’, are present at low numbers within the Yp probe sequence.

**Table 3.**
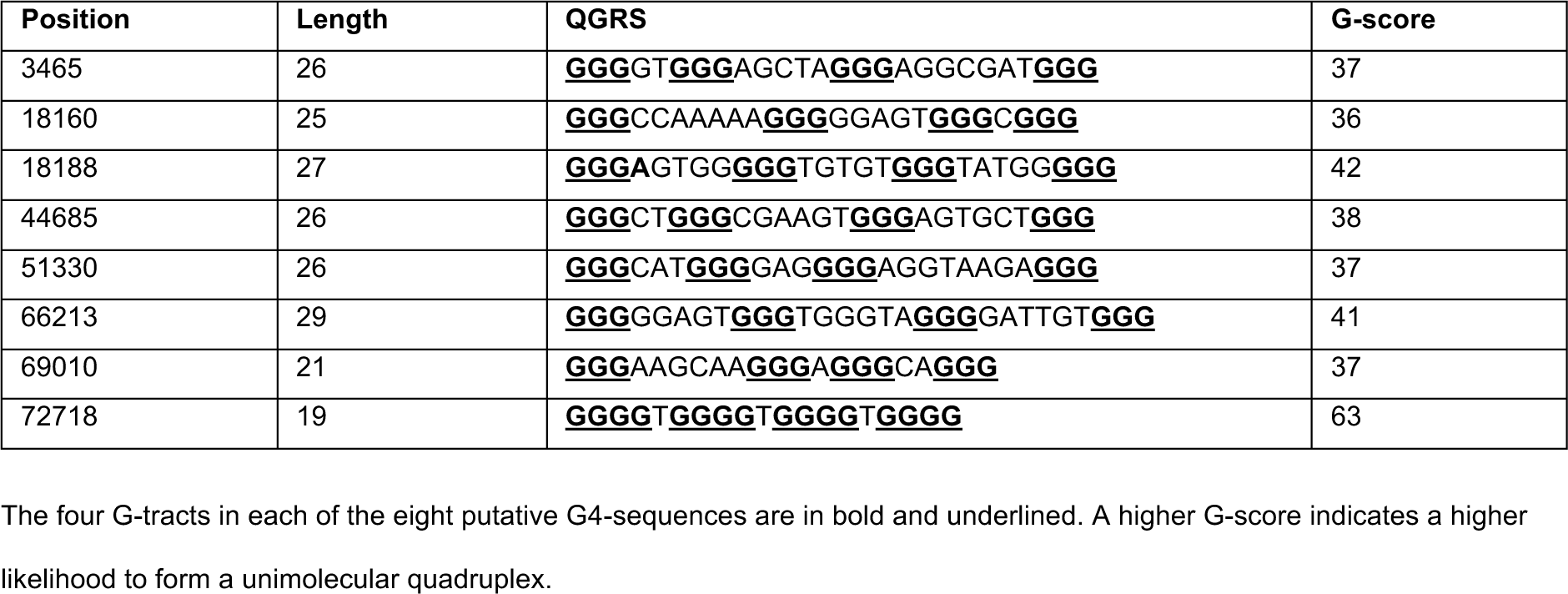
Identification of putative G4-sequences within the Yp probe using QGRS Mapper.

## DISCUSSION

In this study, we have identified a vulnerability of Sertoli cells during fetal development. We show that loss of ATRX in mouse Sertoli cells results in severe local defects, DNA double strand breaks (DSBs) and loss of the chromatin marks HP1α and PH3, at a novel giant GATA4 PML NB on the short arm of the Y chromosome (Yp). In contrast, the vast majority of studies in ATRX-deficient mouse cell lines or tissues such as the brain, mouse ES cells, and oocytes showed that defects like DSBs at telomeres or loss of PH3 at pericentromeric heterochromatin occur across all chromosomes (Baumann et al., 2010; Watson et al., 2013; Wong et al., 2010). The reason is that the chromosomal sites, the telomeres and pericentromeric heterochromatin, where ATRX binds to and is important in these other tissues, are highly conserved in mouse chromosomes (Guenatri et al., 2004). Therefore, loss of ATRX in these tissues will lead to chromosome-wide defects. That in mice a specific PML NB can directly interact with Yp was recently demonstrated in mouse embryonic stem cells (mESC) (Kurihara et al., 2020). In these cells, a 300 kb region (YS300) near the Y chromosome telomere that contains the Yp probe used in our study is frequently associated with a large PML NB (Kurihara et al., 2020). This PML NB forms a specific complex with YS300 to regulate the expression of neighbouring clustered genes such as *Uty* and *Ddx3y*. However, the Yp-linked PML NB in Sertoli cells must serve a different function because these Y-linked genes are spermatogenesis genes and thus are not expressed in Sertoli cells.

Why does a GATA4 PML NB form on Yp in Sertoli cells? All GATA4 foci in control Sertoli cells contain ATRX, DAXX, GATA4 and FOG2, whereas only a subset of GATA4 foci contain HP1α (79%) and PML (17%). This indicates that ATRX, DAXX, GATA4 and FOG2 form a core unit on chromatin which is followed by the recruitment of HP1α and PML. PML is recruited during G2 since almost all the giant GATA4 foci in G2/M-arrested Sertoli cells were positive for PML. DAXX can target ATRX to PML NBs (Ishov et al., 2004; Tang et al., 2004), and *vice versa*, we observed that ATRX is required for expression of DAXX at the PML NBs. Moreover, we found that in Sertoli cells, ATRX is required for localization of HP1α to PML NBs. This is consistent with a previous observation during neural differentiation where ATRX plays a role in targeting HP1α to pericentromeric heterochromatin (Marano et al., 2019). ATRX itself might bind to the potential G4 DNA sequences present within Yp. Since PML interacts with and can recruit DAXX to PML NBs (Ishov et al., 1999), DAXX might also be involved in the recruitment of PML to the GATA4 foci. However, we speculate that the GATA4 foci play a unique role in PML NB formation specifically at Yp. GATA4 is highly expressed throughout the nucleus and is a DNA sequence specific transcription factor with important functions in testis and heart development (Manuylov et al., 2011; Rajagopal et al., 2007; Tevosian et al., 2002). It was therefore surprising to identify an intense focal accumulation of GATA4 within the nucleus. GATA4 has never been reported to be present within PML NBs (Barroso-Gomila et al., 2021; Dellaire et al., 2003; Van Damme et al., 2010). Similarly, its partner protein FOG2 represents a novel protein within PML NBs. One unique feature of the short arm of the mouse Y chromosome is the presence of the evolutionarily conserved banded krait minor (Bkm) satellite DNA sequences in which GATA repeats are the major component (Singh & Jones, 1982; Singh et al., 1994). Indeed, we found that the 73.6 kb Yp probe contains a ∼4.6 kb region with a high accumulation of these GATA repeats. GATA4 could bind repetitively to this region and act as a docking site for PML NB formation. In support of the latter point, GATA2, another member of the GATA family of transcription factors, is one of over 250 proteins associated with PML NBs (Van Damme et al., 2010). Since GATA2 can interact with the PML protein via its zinc finger region shared by all GATA proteins (Tsuzuki et al., 2000), GATA4 may also interact with PML. Another unique feature of Yp is the presence of specific inverted repeats within YS300 that are not homologous to other sequences in the genome (Kurihara et al., 2020). It is therefore possible that these sequences may also play a unique role in the formation of a PML NB specifically at this chromosomal region. That ATRX can associate with specific regions on the mouse Y chromosome was previously demonstrated in male mouse primary embryonic fibroblasts (MEFs), in which ATRX is found at pericentromeric repeat sequences specific to the Y chromosome (Baumann et al., 2008).

One major defect in Sc*Atrx*KO Sertoli cells is the formation of double-stranded breaks (DSBs), as indicated by the presence of γ-H2AX and 53BP1 at a specific HP1α-positive heterochromatic region on Yp that is associated with the GATA4 PML NB. DSBs result from replication stress at chromosomal regions that are difficult to replicate, including common fragile sites, pericentromeric heterochromatin, and telomeres, to which ATRX all binds (Law et al., 2010; McDowell et al., 1999; Pladevall-Morera et al., 2019). The expanded repeats in common fragile sites and telomeres form DNA secondary structures such as hairpins (at common fragile sites) and G4 structures (at common fragile sites and telomeres) that can lead to replication fork stalling or collapse with consequent generation of DSBs (Wang & Patel, 1992; Zlotorynski et al., 2003). ATRX is a crucial factor in common fragile site, pericentromeric heterochromatin, and telomere stability. ATRX facilitates replication by preventing G4 structure formation and promotes replication fork recovery and HR-dependent repair of DSBs (Clynes et al., 2014; Juhász et al., 2018; Leung et al., 2013; Pladevall-Morera et al., 2019; Raghunandan et al., 2020; Watson et al., 2013; Wong et al., 2010). Therefore, one major role of ATRX at the GATA4 PML NB could be to protect a specific HP1α-positive heterochromatic region on Yp that is vulnerable to DNA damage during replication. However, it remains unknown what makes this region on Yp so fragile. The Yp probe sequence contains several putative G4-sequences, therefore, like at telomeres, ATRX may be involved in preventing G4 DNA structures. It is also possible that the GATA repeats in the Yp probe represent a common fragile site that ATRX helps to stabilize. In support of this, a major class of common fragile sites are interrupted runs of AT-dinucleotides that can form secondary structures (Zlotorynski et al., 2003). Of interest, the DNA damage (single γ-H2AX foci) in Sc*Atrx*KO Sertoli cells at the single GATA4 foci often occurs close to the nuclear periphery analogous to mice with conditional knockout of *Atrx* in the limb mesenchyme which also show single γ-H2AX foci close to the nuclear lamina, although those mechanisms remain elusive (Solomon et al., 2013).

A second major defect at the GATA PML NBs in Sertoli cells of Sc*Atrx*KO mice is the loss of HP1α and PH3, indicating changes in chromatin structure and abnormal chromosome condensation, respectively. The GATA4 foci show some structural and functional similarities to chromocenters. Chromocenters are the bright DAPI regions in the nucleus that contain the pericentromeric heterochromatin of several chromosomes (Guenatri et al., 2004). Pericentromeric heterochromatin which is composed of major satellite DNA represent the initiation sites for chromosome condensation, marked by early phosphorylation of histone H3 during G2 (Hendzel et al., 1997). ATRX locates to pericentromeric heterochromatin (McDowell et al., 1999) and mediates its establishment and maintenance through deposition of H3.3, together with DAXX within PML NBs, and recruitment of HP1α (Drané et al., 2010; Marano et al., 2019). Moreover, in mouse oocytes, loss of ATRX leads to reduced phosphorylation of histone H3 at pericentromeric heterochromatin associated with incomplete chromosome condensation and centromeric breaks (Baumann et al., 2010). In Sertoli cells, we found that ATRX is expressed at the HP1α-positive pericentromeric heterochromatin as expected, but also strongly at the HP1α-positive heterochromatin region at the GATA4 foci on Yp. Strikingly, this region is often in very close association with one of the chromocenters. Like the chromocenters, this region contains ATRX, DAXX, HP1α and PML. Moreover, although the Y chromosome lacks the major satellite DNA of the autosomes (Pardue & Gall, 1970), the HP1α-positive Yp heterochromatin region also shows early phosphorylation of histone H3 during G2. Therefore, the GATA4 foci may represent a starting point for Yp-chromosome condensation. Alternatively, early enrichment of PH3 is because heterochromatin in general is condensing early (Drouin et al., 1991). Despite the reported roles for ATRX at pericentromeric heterochromatin, prominent γ-H2AX staining and loss of HP1α and PH3 in Sc*Atrx*KO mice were only observed at the GATA4 foci and not at the chromocenters. This underscores the multiple roles ATRX plays during development, the importance of which depends on the specific tissue context.

The chromatin defects in the G2/M arrested Sertoli cells of Sc*Atrx*KO mice are associated with the occurrence of single giant PML NBs. Intriguingly, single giant PML NBs generated during the G2 phase have been reported once before in a genetic disease, the <u>i</u>mmunodeficiency, <u>c</u>entromeric region instability, <u>f</u>acial anomalies syndrome (ICF) caused by mutations in the DNA methyltransferase gene *DNMT3B* (Ehrlich et al., 2006; Luciani et al., 2005; Xu et al., 1999). DNMT3B is involved in the methylation of GC-rich major satellites that are predominantly located at the pericentromeric heterochromatin of chromosomes 1 and 16 (satellite 2 DNA) and chromosome 9 (satellite 3 DNA). *DNMT3B* mutations lead to hypomethylation, decondensation and instability of these major satellites, resulting in chromosome breaks. The giant PML NBs in ICF cells consist of ATRX, DAXX, HP1α and several other proteins and contain the major satellites of all three chromosomes. It was therefore proposed that these PML NBs promote the condensation of these satellites specifically at G2 phase before mitosis. The authors also concluded that the giant PML NBs in ICF syndrome patients are most likely due to the undercondensed satellite DNA occupying a larger space compared to the condensed state. Given these data and the absence of HP1α and PH3 at the giant GATA4 PML NBs in Sc*Atrx*KO Sertoli cells, it is likely that the Yp region occupied by the giant PML NB is also undercondensed. Overall, these data indicate that the PML NB associated with chromosomes 1, 9 and 16 in humans and the PML NB associated with the Y chromosome in mice show functional similarities. Therefore, a major role of ATRX at the GATA4 PML NB may be to maintain the heterochromatic state and to promote condensation of the HP1α-positive heterochromatic region on Yp. In mice, the association of a PML NB to the pericentromeric heterochromatin of specific autosomes appears unlikely, because the major satellites are highly conserved between chromosomes except for the Y chromosome (Guenatri et al., 2004)

The occurrence of two major defects at the GATA4 PML NBs of Sc*Atrx*KO mice, DSB formation and a failure of chromosome condensation, could reflect distinct roles for ATRX in Sertoli cells. However, it is also possible that ATRX is not directly required for recruitment of HP1α and phosphorylation of histone H3 at the GATA4 foci and that the chromatin structure and condensation defects are a consequence of the DNA double strand breaks. Indeed, DNA damage response can result in dynamic changes in chromatin structure, for example experimental induction of double stranded breaks in mammalian cells leads to local chromatin decondensation (Berkovich et al., 2007; Kruhlak et al., 2006; Ziv et al., 2006).

Based on our findings, we propose that in Sc*Atrx*KO mice, DSBs are generated at a fragile Yp heterochromatic region within a dysfunctional PML NB. This will trigger the phosphorylation of the Histone variant H2AX (γ-H2AX) which is known to recruit DNA repair proteins and induce G2/M cell cycle arrest (Plesca et al., 2008; Rogakou et al., 2000). Either as an independent event within the dysfunctional PML NB or due to the DSBs, the heterochromatic state of the Yp region is not maintained which fails to condense, resulting in an expansion of the PML NB (giant PML NB). All Sertoli cells with giant PML NBs were arrested at G2/M and showed γ-H2AX-staining on their giant PML NBs, about half of them also showed widespread γ-H2AX-staining. These observations directly link chromosome decondensation and DSBs to the observed G2/M arrest and apoptosis in Sc*Atrx*KO mice.

In summary, our data indicate that DNA damage and a failure of chromosome condensation at the short arm of the Y chromosome underly the testicular abnormalities in ATR-X syndrome. This establishes Y chromosome damage as a novel mechanism for testicular failure.

## MATERIALS AND METHODS

### Mouse generation and genotyping

*Atrx^flox/flox^* female mice were crossed with *AMH-Cre/+* male mice to generate *AMH-Cre*/+;*Atrx^flox/Y^* (Sc*Atrx*KO) male mice. *Atrx^flox/Y^* male mice were used as controls. For embryonic time points, noon of the day after mating was considered E0.5. Dissection of whole embryos was performed at E16.5 and E17.5. For genotyping, genomic DNA was isolated from tail tissue and subjected to genotyping using a previously published protocol (Barrionuevo et al., 2006; Garrick et al., 2006). All animal experimentation was approved and performed according to procedures determined by the Monash Medical Centre Animal Ethics Committee.

### Tissue processing and paraffin sectioning

Mouse embryos were collected at day E16.5 and E17.5, washed with 1x PBS and fixed with 4% paraformaldehyde (PFA). Fixed embryos were sent to the Histopathology platform at the Hudson Institute of Medical Research for tissue processing and paraffin embedding. Sagittal sections of the embryos were generated with a microtome at 4μm thickness.

### Immunofluorescence

Paraffin testis sections of Sc*Atrx*KO and control *Atrx^flox/Y^* male mice were sent to the Histopathology platform at the Hudson Institute of Medical Research for dewaxing. The slides were heated at 60° degrees and dipped in xylene to remove the paraffin. Then, the slides were placed in 100% EtOH 3 times for 3 minutes each followed by dipping in distilled water (dH2O) for 5 minutes. Heat-induced epitope retrieval was done using a pressure cooker containing 15 ml of Antigen Citrate-based unmasking solution (Vector laboratories; H-3300), diluted in 1.6 L of dH2O. For immunofluorescence, sections were blocked in blocking solution (1x PBS with 0.1% Tween20 and 5% donkey serum) for 1 hour at room temperature. Then, the sections were incubated with diluted primary antibodies at 4°C overnight. On the next day, the slides were washed with 1X PBS 3 times for 5 minutes each and incubated with secondary antibodies for 1 hour. Table S1 contains a list of the primary and secondary antibodies used in this study.

To reduce autofluorescence, Sudan Black solution (0.1% in 70% EtOH) was applied to the slides which were incubated at room temperature for 8 minutes. After washing with 1X PBS, the slides were counterstained with DAPI (1:5000) for 5 minutes at room temperature, and then washed in 1X PBS. Finally, the slides were mounted using fluorescent mounting medium (Dako). Imaging was performed using confocal microscopes (Olympus FV1200 and Nikon C1, Monash Imaging).

### Preparation of BAC DNA

The BAC clone B6Ng01-016L17 which contains 73.6 kb of the short arm of the Y chromosome was provided by Dr. Yusuke Miyanari from Kanazawa University. For preparation of the BAC DNA, the BAC strain was first cultured on a LB agar plate containing 25μg/ml of chloramphenicol and incubated at 37°C overnight. Then, a single colony was cultured in 2 ml of LB Broth media with 25μg/ml of chloramphenicol and incubated at 37°C for 8 hours. The 2 ml were added to 500 ml of LB Broth media with 25 μg/ml of chloramphenicol and incubated overnight at 37°C. The culture medium was then centrifuged at 4,000 rpm for 10 minutes at room temperature. The BAC DNA was purified using the NucleoBond Xtra BAC kit (Macherey-Nagel 740436.2) following the manufacturer’s protocol. Finally, BAC DNA was resuspended in 700 μl of sterile H_2_O and incubated overnight at 4°C to dissolve completely. The sequence of the BAC DNA was confirmed by Sanger sequencing using the primers listed in Table S2.

### Immuno-FISH

Dewaxed sections of Sc*Atrx*KO and control *Atrx^flox/Y^* mice were first stained with a GATA4 antibody using the immunofluorescence protocol described above. Afterwards, FISH was performed starting with the dehydration of the slides in an ethanol series of 75%, 80% and 100%, and then air-dried. To denature, slides were placed in 2X SSC/Formamide at 80°C for 5 minutes. After drying the slides, they were treated with a cold ethanol series of 75%, 80%, and 100%, and then dried again. The Yp and Yq DNA probes (Table S3) were heated at 95°C for 5 mins and immediately placed on ice.

The probe was added to the slide and a coverslip was placed on top, sealing the edges with glue. The slides were then placed in an opaque and humidified chamber and were incubated at 37°C overnight. The next day, the slides were washed 3 times with 2X SSC for 5 minutes each, followed by 2 times with 0.5X SSC for 5 minutes each, at room temperature. TNB block was added to each slide and incubated for one hour at 37°C. Next, the slides were incubated with AV488 (1/400) for one hour at 37°C. The slides were then washed 3 times with 4X SSC + 0.1% Tween-20 for 5 minutes each time. They were quickly rinsed in PBS solution, fixed in 4% PFA for 7 minutes and air-dried. Finally, for nuclei detection, DAPI was applied, and the slides were kept in the dark at 4°C. Imaging was performed using confocal microscopes (Olympus FV1200 and Nikon C1, Monash Imaging).

### Quantification and statistical analysis

#### Quantification of cell numbers

To determine the average percentage of Sertoli cells that were positive for GATA4, PH3, γ-H2AX and PML, the total number of Sertoli cells and marker-positive cells was manually counted, using cell count tool of FUJI software. More than 800 Sertoli cells were analysed in each testis (control n=2; Sc*Atrx*KO=3). A second blind observer also performed the quantification, obtaining similar results. Graphs and statistical analysis were done using GraphPad Prism v8.1.

#### Measurement of GATA4 foci diameter

A total of 3 control and 4 Sc*Atrx*KO testes were analysed, using FUJI software. More than 100 GATA4 foci and GATA4-PML NBs were measured in each testis. To determine their size a threshold was applied as the GATA4 foci/ GATA4-PML NBs presented the most intense signal in the image. Then, specific parameters were set based on the scale bar of each image, and the program did an automatic measurement of the area and perimeter of the foci. All images were analysed under the same parameters. Finally, GraphPad Prism v8.1 software was used to obtain graphs and perform the statistical analysis.

All statistical analysis was conducted with GraphPad Prism v8.1, using appropriate tests as stated in the figure legends. Statistical significance was determined by two-tailed t-test or one-way ANOVA analyses. A p-value <0.05 was utilized as the cut-off for statistical significance.

## Acknowledgements

We thank Monash Medical Imaging, Monash Histology, Helena Sim, Janelle Ryan, Sophia Astbury, Kerryn Bird and Karisma Lawrence for technical assistance, and Florian Guillou for *AMH-Cre* mice and Doug Higgs for the *Atrx^floxflox^* mice.

---

**Figure S1. GATA4 and strong ATRX labelling co-localize in Sertoli cells near a highly compacted heterochromatin region.**

Double immunofluorescence (IF) analyses in E17.5 XY control and Sc*Atrx*KO testes for GATA4 (red, nuclear) and ATRX (green, nuclear). DAPI (blue) was used as a nuclear stain. White arrows denote ATRX-positive pericentromeric heterochromatin (bright DAPI regions), and yellow arrows show co-localization of strong ATRX staining with the GATA4 foci. Scale bar is 5μm.

**Figure S2. Sertoli cells show DNA double strand breaks at the GATA4 foci.**

Double immunofluorescence (IF) analyses in E16.5 XY control and Sc*Atrx*KO testes for GATA4 (red, nuclear) and 53BP1 (green, nuclear). White arrows denote GATA4 foci. Scale bar is 5μm.

